# Investigating speciation in face of polyploidization: what can we learn from approximate Bayesian computation approach?

**DOI:** 10.1101/002527

**Authors:** Camille Roux, John R. Pannell

## Abstract

Despite its importance in the diversification of many eucaryote clades, particularly plants, detailed genomic analysis of polyploid species is still in its infancy, with published analysis of only a handful of model species to date. Fundamental questions concerning the origin of polyploid lineages (e.g., auto- vs. allopolyploidy) and the extent to which polyploid genomes display disomic vs. polysomic vs. heterosomic inheritance are poorly resolved for most polyploids, not least because they have hitherto required detailed karyotypic analysis or the analysis of allele segregation at multiple loci in pedigrees or artificial crosses, which are often not practical for non-model species. However, the increasing availability of sequence data for non-model species now presents an opportunity to apply established approaches for the evolutionary analysis of genomic data to polyploid species complexes. Here, we ask whether approximate Bayesian computation (ABC), applied to sequence data produced by next-generation sequencing technologies from polyploid taxa, allows correct inference of the evolutionary and demographic history of polyploid lineages and their close relatives. We use simulations to investigate how the number of sampled individuals, the number of surveyed loci and their length affect the accuracy and precision of evolutionary and demographic inferences by ABC, including the mode of polyploidisation, mode of inheritance of polyploid taxa, the relative timing of genome duplication and speciation, and effective populations sizes of contributing lineages. We also apply the ABC framework we develop to sequence data from diploid and polyploidy species of the plant genus *Capsella*, for which we infer an allopolyploid origin for tetra *C. bursa-pastoris* ≈ 90,000 years ago. In general, our results indicate that ABC is a promising and powerful method for uncovering the origin and subsequent evolution of polyploid species.

## INTRODUCTION

Our ability to use of contemporary patterns of genetic variation to infer and understand past evolutionary and demographic processes (Harrison 1993) has taken great strides in recent years with the availability of inexpensive sequence data and new statistical tools to analyze it. One of the aims of such projects is to infer the demographic history of species from patterns of polymorphism and divergence observed within and among natural populations. The rationale beyond this approach is that past events, such as a strong bottleneck within a lineage (Gattepaille et al. 2013) or a secondary contact between divergent populations (Durand et al. 2011), leave distinct genetic signatures in gene pools, which can be used to infer the nature and timing of the processes that gave rise to them. For example, after the occurrence of a severe bottleneck, linkage disequilibrium (LD) extends to larger genomic regions than one would expect to find in stable populations (Reich et al. 2001), coalescent trees became star-shaped, and there is a resulting excess of rare mutations across the whole genome. Such patterns of genetic variation have allowed the inference of bottlenecks and subsequent population expansion in recent history of a number of species, including *Drosophila melanogaster* (Li and Stephan 2006), *Arabidopsis thaliana* (François et al. 2008) and *Oryza sativa* (Caicedo et al. 2007).

The rise of approximate Bayesian computation (ABC) (Tavaré et al. 1997; Pritchard et al. 1999; Beaumont et al. 2002) has made a substantial impact on our ability to infer the demographic and evolutionary history of populations from genetic data. ABC is a tool for statistical comparisons between alternative models of evolution (Beaumont et al. 2010). It avoids the difficulty of computing the likelihood function of complex demographic models by comparing the values of a vector of summary statistics between an observed genetic dataset data produced by simulations. By using a rejection-and-regression algorithm (Beaumont et al. 2002; Blum and François 2009), the posterior probability of a model *M* can then be approximated by its relative acceptance frequency.

ABC has successfully been used to infer the demographic history and history of evolutionary divergence for a wide range of organisms. For instance, comparing different scenarios of human evolution using ABC, Fagundes et al. (2007) provided unambiguous support for a recent human colonization out of Africa about 50,000 years ago, with no long-term gene flow between continents, thus contributing to the rejection of the ‘multiregional hypothesis’ (Cavalli-Sforza and Feldman 2003; Lahr 1994; Mellars and Stringer 1989). To understand the history of divergence in the plant genus *Arabidopsis*, Roux et al. (2011) used ABC to reveal a tight relationship between the establishment of a major adaptive mutation in *Arabidopsis* and speciation. Inferring realistic demographic models from genetic data leads to better knowledges about the history of species, but it is also crucial for building statistical null models for the detection of molecular signatures of direct and indirect selection across genomes (Nielsen 2005; Roux et al. 2012).

Coalescent-based ABC has hitherto been mainly applied to haploid and diploid organisms, with little attention given to polyploid species, despite their recognized importance (Jakobsson et al. 2006; St Onge et al. 2012; Lepais et al. 2013). Indeed, polyploidy has played a fundamental role in the evolution and diversification of a wide range of taxa, including invertebrate and vertebrate animals and, especially, plants (Otto and Whitton 2000; Gregory and Mable 2005; Gallardo et al. 2004; Neiman et al. 2011). Concerning plants, the estimated proportion of current species with more than two copies of each chromosome is claimed to reach values as high as 35% (Arrigo and Barker 2012), with its significance having been revealed at very different timescales. At one extreme, for example, comparative genome analysis has revealed a paleopolyploidization event that predates the radiation of angiosperms (Bowers et al. 2003); at the other extreme, several new polyploid species have been reported to have become established in the past 150 years (Soltis et al. 2003). Within a lineage, polyploidization has been proposed as a mechanism of species differentiation by favoring instant sympatric speciation because of the low fitness or sterility of hybrids between diploid parents and their polyploid descendants (Coyne et al. 2004; Wood et al. 2009; Linder and Rieseberg 2004; Mallet 2007). Consistent with this mechanistic, Mayrose et al. (2011) compared speciation rates between diploids and neo-polyploids in angiosperms and seed-free vascular plants and confirmed that whole-genome duplication (WGD) of diploids has been a major process driving speciation, albeit counterbalanced to some extent by a greater extinction rate of neopolyploids over diploids (Mayrose et al. 2011).

Despite the importance of polyploidization in speciation, significant theoretical and empirical approaches to fit polyploid genomic data to evolutionary or demographic models have emerged only recently (Jones et al. 2013). A pivotal issue concerns determining the effects of genome duplication on the coalescent model (Hudson 1983; Tajima 1983) in order to delimit what we can do with current tools (Hudson 2002; Laval and Excoffier 2004). In this context, a key question concerns how polyploidization will influence demographic inference within a lineage that is likely to have gone through one or more genetic bottlenecks, that may have evolved more than once, and for which (in the case of allopolyploidy) the ancestral population was subdivided into two potentially reproductively isolated units. Not only do these different scenarios potentially complicate demographic inference; they are also intrinsically interesting in themselves. How can recent advances in demographic and evolutionary inference help us to infer the nature and timing of these different events?

One is often interested in distinguishing between the establishment of a polyploid lineage following the simple doubling of the genome within a single population (autopolyploidy) and its establishment as a result of hybridization and genome doubling between two divergent populations or species (allopolyploidy). To model autotetraploid populations, Arnold et al. (2012) recently developed a coalescent framework that assumes tetrasomic inheritance (so that all of the six possible gametes have the same probability of being produced by tetraploid adults) and showed that a rescaled Kingman model for haploids serves as a suitable approximation. Lineages produced by allopolyploidization of course have a more complex history, not least because they combine two genomes that can be potentially quite divergent, each originating from a different species with its own demographic history. The demographic history of an allopolyploid genome thus includes three different histories: the two histories of the contributing parental species, and the history following hybridization and effective genome duplication. Demographic inference involving allopolyploid lineages thus needs to account for, and estimate, the histories of each of these component populations.

There have been substantial recent advances in our understanding of the rôles played by auto- versus allopolyploidisation in speciation between diploid and polyploid species (Jakobsson et al. 2006; Slotte et al. 2008; Wang et al. 2010; St Onge et al. 2012). For instance, St. Onge et al. (2012) used the MIMAR software (Becquet and Przeworski 2007) to distinguish between auto- and allopolyploidization in the divergence between diploid plant *Capsella grandiflora* and its tetraploid derivative *C. bursa*-pastoris, estimating the parameters of the Isolation with Migration (IM) model. In their analysis, they considered both sub-genomes A and B of *C. bursa-pastoris* as effectively independent species experiencing divergent evolution, and compared them with the genome of *C. grandiflora*. They then estimated speciation times between *C. grandiflora* and the *C. bursa-pastoris* A and B genomes, respectively. Although this approach should be straightforward, it is not compatible with modern datasets because it requires the specific assignment of each of the four phased-haplotypes to one of the two diploid sub-genomes (homeologous pairs) for the tetraploid species (but see (Salmon et al. 2012)). MIMAR was originally designed to estimate parameters of the IM model only (the effective population sizes of the ancestral population *N*_A_ and for both daughter populations *N*_1_ and *N*_2_, the time of speciation *T*_split_, and migration rates in both directions), and it does not provide an explicit statistical test of alternative scenarios of speciation with polyploidization.

Although the distinction between auto- and allopolyploidy is conceptually useful, it is important to note that notably the distinction between tetrasomic and disomic inheritance between the two modes of polyploidisation are expected to be transitory. Typically, allopolyploid lineages will show disomic inheritance, with preferential pairings between the two homologous chromosomes of each contributing species, i.e., between the homeologues; in this case, the two sub-genomes of an allotetraploid lineage can be seen as two completely isolated diploid populations, because recombination occurs only between the chromosomes contributed by each parent species, and not between them. In contrast, genome duplication leading to autotetraploidy will typically give rise to a lineage with initially tetrasomic inheritance, in which homologous chromosomes will pair at random, and recombination can occur between any of the four chromosome. But tetrasomic inheritance in a recently duplicated tetraploid lineage will gradually give way to disomic inheritance as the polyploid genome becomes progressively diploidised as a result of the divergence between two different homeologous chromosome pairs. Because this process of diploidisation can be prevented at islands of recombination between the two sub-genomes (Meirmans and Van Tienderen 2013), differences in gene exchange between homeologous pairs across the genome can lead to heterosomy, with some loci showing disomic inheritance and others showing tetrasomic inheritance. Heterosomy has somewhat neglected in the literature (Allendorf and Danzmann 1997; Koning-Boucoiran et al. 2012; Stift et al. 2008; Justin and W. 2002), but it should have profound consequences on the distribution of coalescent times across a genome and thus on demographic and evolutionary inference.

In addition to the mode of origin of a polyploid lineage, an important question concerns the relative timing genome duplication and divergence from its diploid progenitor(s): e.g., does speciation precede genome duplication, or do the two processes coincide (Fig. 1)? A striking observation in plants (Obbard et al. 2006) and animals (Neiman et al. 2011) is that the transition from diploidy to tetraploidy is frequently associated with a shift from obligate outcrossing to predominant self-fertilization (Astaurov 1969; Otto and Whitton 2000). One hypothesis is that this pattern results from the breakdown of self-incompatibility with polyploidization, which the subsequent divergence of the new tetraploid lineage. Alternatively, the subdivision of an ancestral diploid species through colonization may lead to the establishment of self-compatible individuals, e.g., through selection for reproductive assurance (Baker 1955), followed by genome duplication; in this case, genome duplication might occur well after the establishment of self-compatibility because of drift, or because of possible selective advantages conferred by tetraploidy (Chao et al. 2013). To what extent can genetic data allow us to distinguish between these two alternatives using an ABC framework?

**Figure 1.**
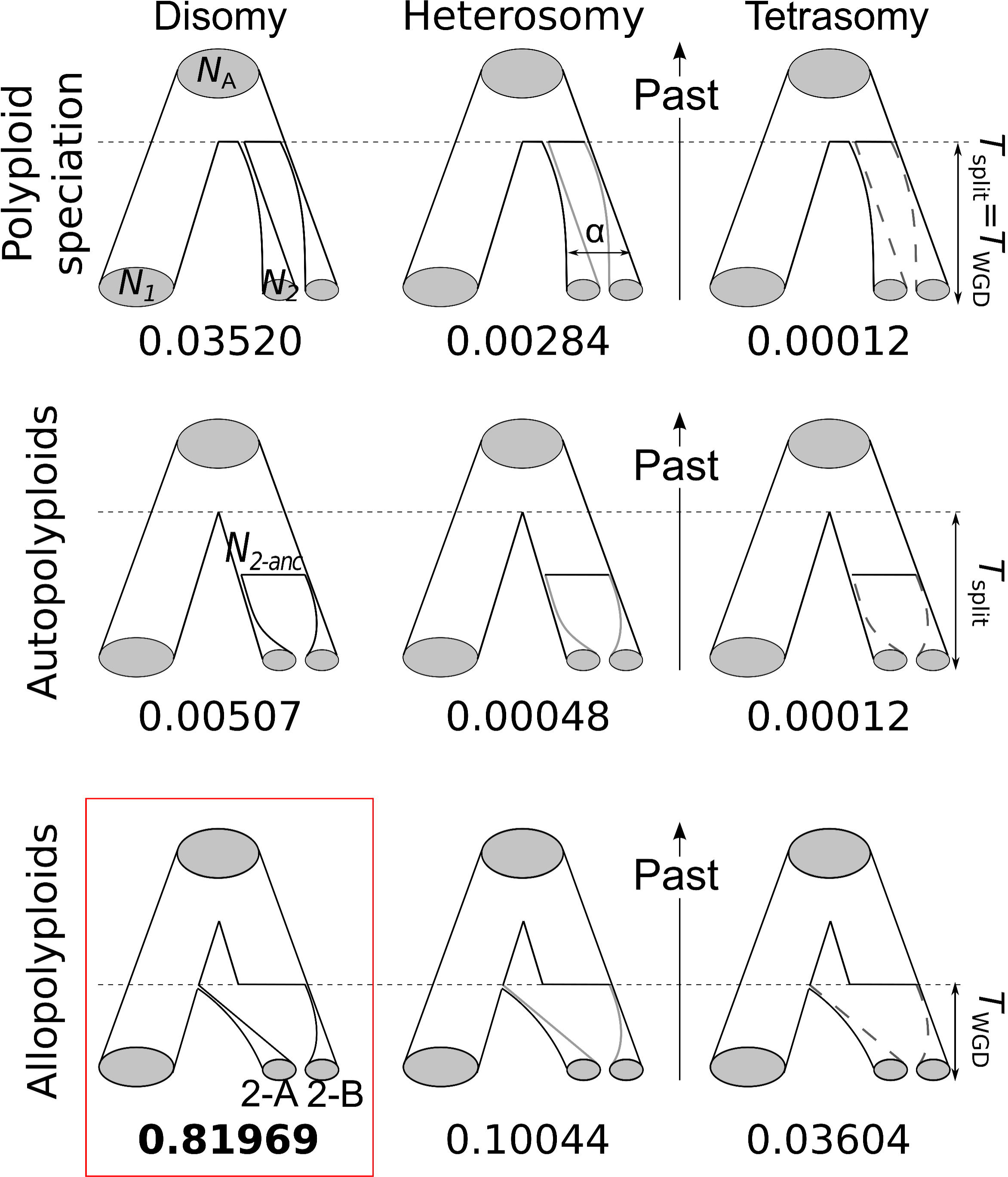
Alternative scenarios of speciation and polyploidization. Parameters are as follows. *N*_1_: effective population size of the current diploid species. *N*_2_: effective population size of each sub-genomes making the tetraploid species. *N*_2A_: effective size of the diploid population ancestral to the tetraploid species, for autopolyploidization and allopolyploidization scenarios only. *N*_A_: effective population size of the common ancestor of diploid and tetraploid species. *T*_split_: time of speciation. *T*_WGD_: time of whole-genome duplication. α: proportion of the genome associated to tetrasomic inheritance, for heterosomic species only. Numbers are the relative posterior probabilities attributed by ABC from the *Capsella* dataset when all models were compared in the same analysis. The red rectangle indicates the best-fitting model to the *Capsella* dataset.

In this article, we evaluate the extent to which ABC modelling allows us to recapture the evolutionary and demographic history of polyploid lineages using data (e.g., generated by next-generation sequencing; NGS) with which it is not possible to know either the haplotypic phase of sequence data or the sub-genome from which it originated. Specifically, we use ABC in an attempt to recapture several of the key processes that might have led to the establishment of a tetraploid lineage from one or more progenitor diploid populations, comparing nine different models (Fig. 1). These models vary in: the mode of origin of polyploidy (polyploid speciation, auto- and allopolyploidy); the relative timing of genome duplication and population divergence; and the modes of inheritance characterizing the tetraploid lineage (disomy, heterosomy or tetrasomy). In addition, we ask both how inferential power might be improved when sequence data from an outgroup are available, and how inference on the basis of a NGS-like dataset (with a large number of loci over few individuals) compares with that based on Sanger sequencing (with fewer loci and more individuals sampled). Finally, we apply our approach to the analysis of polymorphism and divergence from published DNA sequences from diploid and tetraploid lineages of the plant genus *Capsella*, for which hypotheses regarding polyploidization and demographic history have been considered using different approaches (Slotte et al. 2006, 2008; St Onge et al. 2012).

## Results and discussion

### Distinguishing between allo- and autopolyploidisation

Our simulations show that ABC can reliably distinguish between an allo- or autopolyploid origin of a tetraploid organism on the basis of a realistic genomic dataset (Fig. 2). We generated 1,000 pseudo-observed datasets for each of the allo- and autopolyploidization models and obtained the distributions of relative posterior probabilities consistent with the evolutionary and demographic model used to generate the sequence data. Importantly, our ability to correctly identify the origin of polyploidization depended to some extent on the mode of inheritance assumed for the sequenced loci. Thus, the proportion of successfully supported pseudo-observed datasets decreased with the intensity of recombination between the two sub-genomes of the derived tetraploid lineage. For instance, a scenario of autopolyploidization was correctly identified in 99%, 93.5% and 83.6% of the simulated cases for disomic, heterosomic and tetrasomic inheritance, respectively. This loss of information with increasing heterosomy can be attributed to recombination between the otherwise homeologous chromosomes during the early stages after polyploidization and recovery from the corresponding population bottleneck caused by the genome duplication of a single individual.

**Figure 2:**
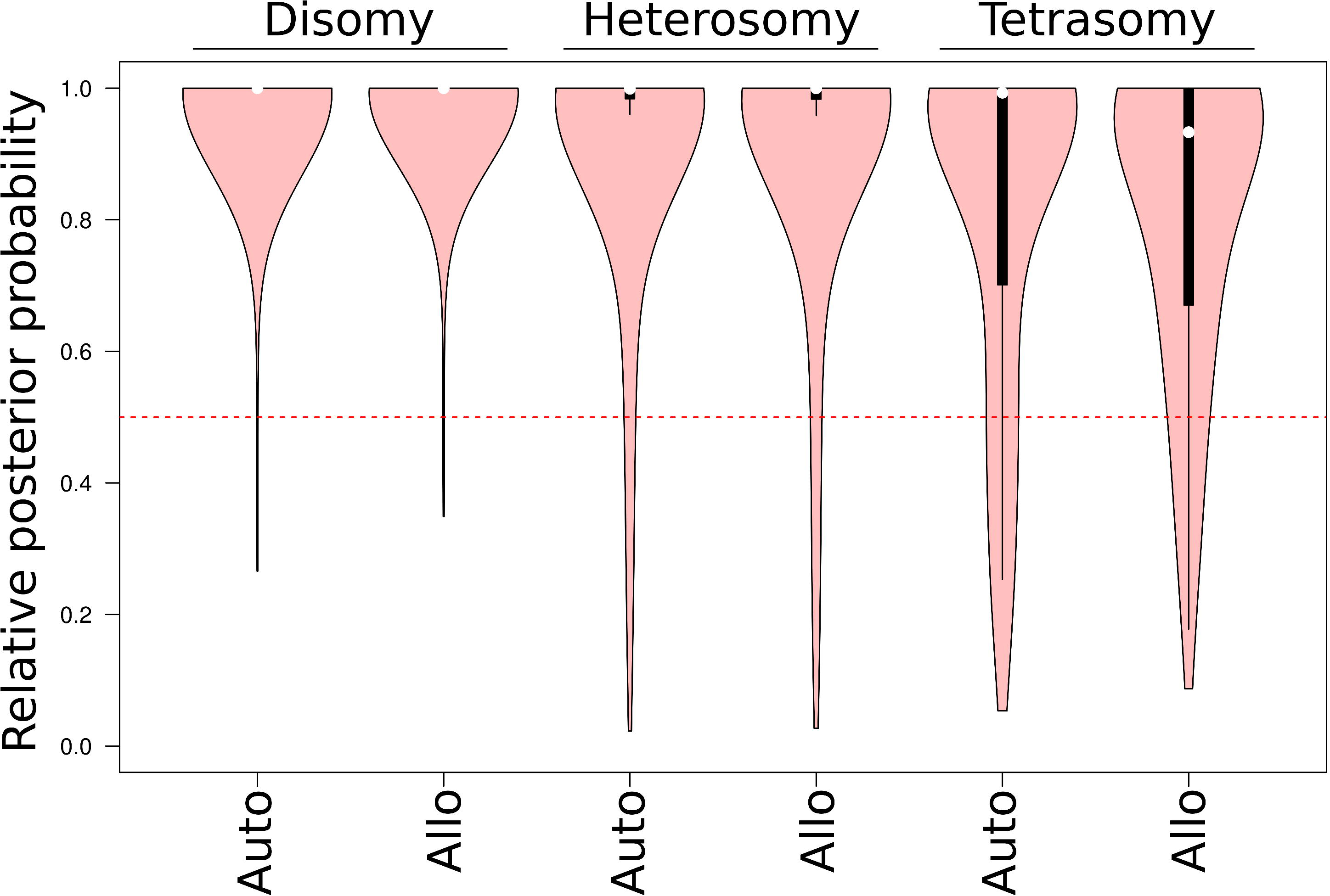
Empirical distributions of relative posterior probabilities to correctly support the polyploid origin regarding the mode of inheritance. Red areas represent the distributions of the estimated values of *P*(Auto | Auto) and *P*(Allo | Allo) obtained after model-comparisons using ABC from 1,000 pseudo-observed datasets randomly simulated under auto- and allopolyploidization scenarios respectively. Pairwise model-comparisons were made for disomic, heterosomic and tetrasomic modes of inheritance. The black boxes within the red areas delimit the 25% and 75% quantiles of each distribution, and the white dots show the median. In pairwise model-comparisons, a model *M* is the best-supported model if its relative posterior probability is greater than 0.5.

Distinguishing between an auto- and allopolyploid origin of a polyploid lineage has hitherto required detailed cytogenetic analysis. Such work included the extraction and staining of metaphasic chromosomes in order to characterize chromosomal configurations, such as the telomeric heterochromatin in the long arm (Tymowska and Fischberg 1982; Chalup et al. 2012), and cluster analysis to test whether each of the homeologous chromosomes group together as expected under autopolyploidy (Gutiérrez et al. 1994). Despite improvements in the acquisition of molecular markers, the rational beyond this test has more or less remained unchanged, with auto- versus allopolyploid origins still distinguished by testing for the occurrence of divergent chromosomes within the same tetraploid cytotype, even where data from parental species are not available (Mráz et al. 2012). The mode of inheritance discerned by cygogenetic analysis is still frequently used to directly infer the polyploid origin (Hardy et al. 2000), even though this relationship may be misleading (Le Comber et al. 2010). Even in recent studies using simulation-based methods, the mode of inheritance is assumed to be disomic by default, so as to infer the origin of polyploidization by way of estimates of divergence dates between each sub-genome pair and putative diploid relatives (St Onge et al. 2012). Our analysis now points to ABC as a powerful means of inference for the origin of polyploid lineages, not least because it is able to account for the demographic history and the diversity in recombinational models between homeologous chromosomes.

### Inferring the mode of inheritance

Our simulations also show that, beyond the fundamental distinction between allo- and autopolyploidisation, it is possible to infer (rather than assume) the mode of inheritance for a polyploid lineages (Fig. 3). Thus, we were able to correctly attribute the mode of inheritance as either disomic, heterosomic or tetrasomic at least 96.9% of the time, regardless of the specific scenario underlying speciation. This result should give us confidence in the statistical support for any model of inheritance provided by ABC on the basis of sequence variation across the genome, without needing to deal with logistically more demanding and complex cytogenetic approaches.

**Figure 3:**
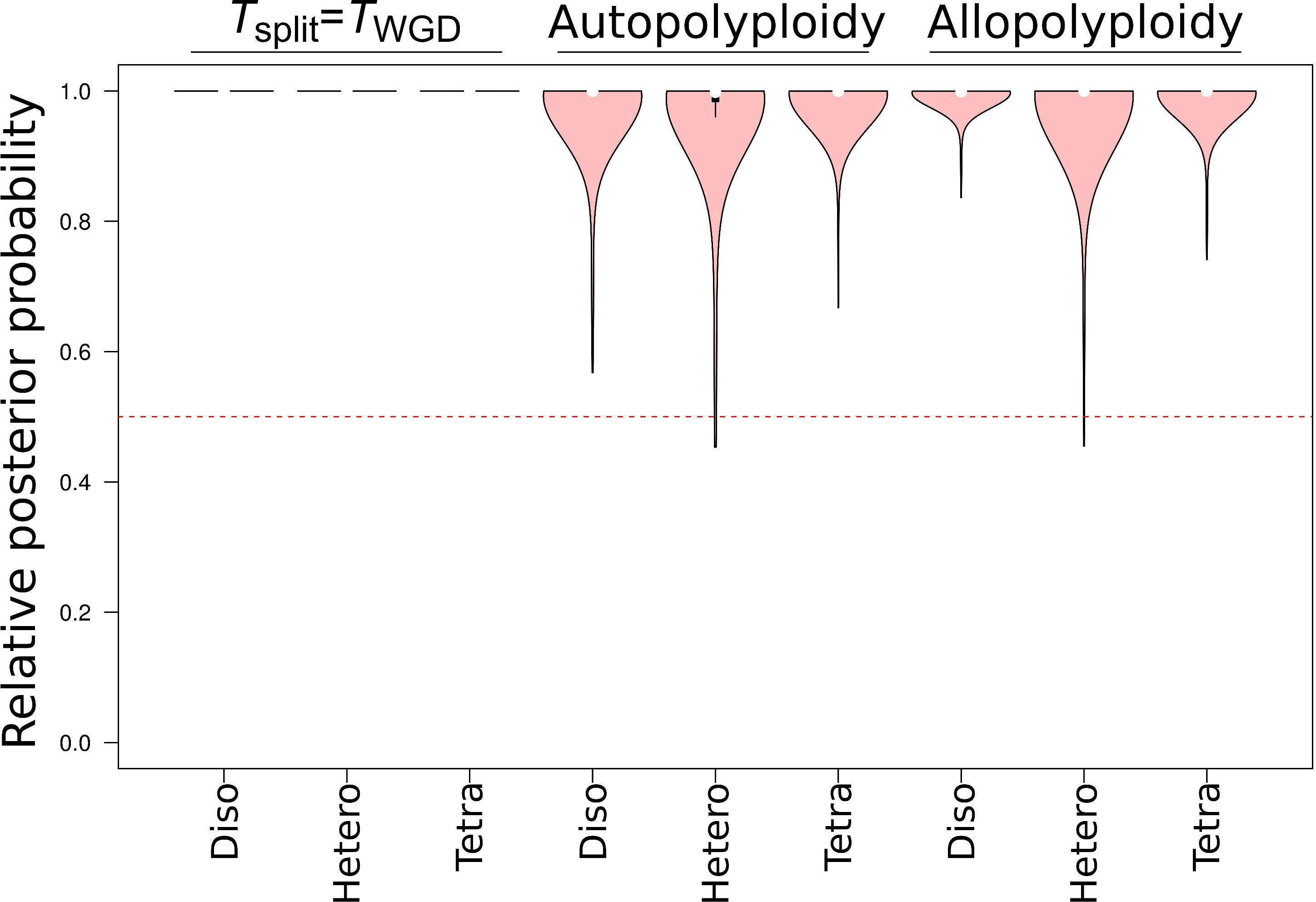
Empirical distributions of relative posterior probabilities to correctly support the mode of inheritance regarding the polyploid origin. Red areas represent the distributions of the estimated values of *P*(Disomy | Disomy), *P*(Heterosomy | Heterosomy) and *P*(Tetrasomy | Tetrasomy) obtained after model-comparisons using ABC from 1,000 pseudo-observed datasets randomly simulated under disomic, heterosomic and tetrasomic modes of inheritance, respectively. Model-comparisons were made for auto- and allopolyploidization scenarios, respectively.

Historically, the approaches to establish the mode of inheritance have been based on intensive cytogenetic studies or segregation analysis from experimental crosses between parents of known genotypes (Kamiri et al. 2011; Pupilli et al. 1997). By using segregation patterns of genetic markers in offspring at multiple loci, is has been necessary to compute the probabilities of the different models of inheritance to finally test the support of the best hypothesis through arbitrary interpretations of the Bayes factor value (Olson 1997). Although recent technological advances have made hundreds of molecular markers available for screening the genomes of non-model species (Gayral et al. 2013), it will often be difficult and/or prohibitively time-consuming to obtain and genotype sufficiently large numbers of progeny for most wild species. Catalàn et al. (2006) have shown that, in some cases, it might be possible to circumvent this problem by making use of microsatellite profiles from natural populations to discriminate between disomic and tetrasomic inheritance without performing crosses, but this approach will rarely be feasible. Some methods have been successfully adopted to accommodate the possibility of intermediate heterosomic models of inheritance, but these, too, make use information from a segregation analysis (Stift et al. 2008) and so will often not be available.

As our study shows, investigating the mode of inheritance can be made using ABC on the basis of DNA sequences, without having to assume a strict relationship between the historical scenario and any biological process. Although we have considered probably the least complex scenarios that one might face, the approach we have used should easily be extended or adapted to accommodate more complex models. A first extension, for example, would be the implementation of a framework to identify a gradual shift from polysomy to disomy, since multivalent pairing is ephemeral (Otto 2007). Such models would provide a better description of the linear or quadratic relationship between the time of polyploidization and the proportion of the genome that has become disomic, but it would also provide a better understanding of the evolutionary forces driving such transitions.

### Testing for the relative timing of duplication and speciation

It is clear from our simulations that ABC can provide a robust test for the co-occurrence of polyploidization and speciation under certain inheritance modes (Fig. 4). We found that ABC is frequently able to recapture the evolutionary and demographic parameters for *T*_split_ = *T*_WGD_ (polyploid speciation) and *T*_split_ > *T*_WGD_ for cases with disomic (with respectively 99% and 96.8% of correctly supported pseudo-observed datasets) and heterosomic inheritance (with respectively 96.8% and 92.4% of correctly supported pseudo-observed datasets). However, with recombination between homoeologous chromosomes, power to recapture the simulated parameters fell dramatically (to values of 58.3% and 55.5% for *T*_split_ = *T*_WGD_ and *T*_split_ > *T*_WGD_, respectively). Under fully tetrasomic inheritance, distinguishing between these two models is similar to estimating the timing of speciation between two undifferentiated gene pools linked by elevated introgression. It is thus clear that a minimum level of preferential pairing between homoeologous chromosomes in the genome will be required for accurately testing for the co-occurrence of polyploidization and speciation. It will thus be important to test for the mode of inheritance prior to attempting such evaluations.

**Figure 4:**
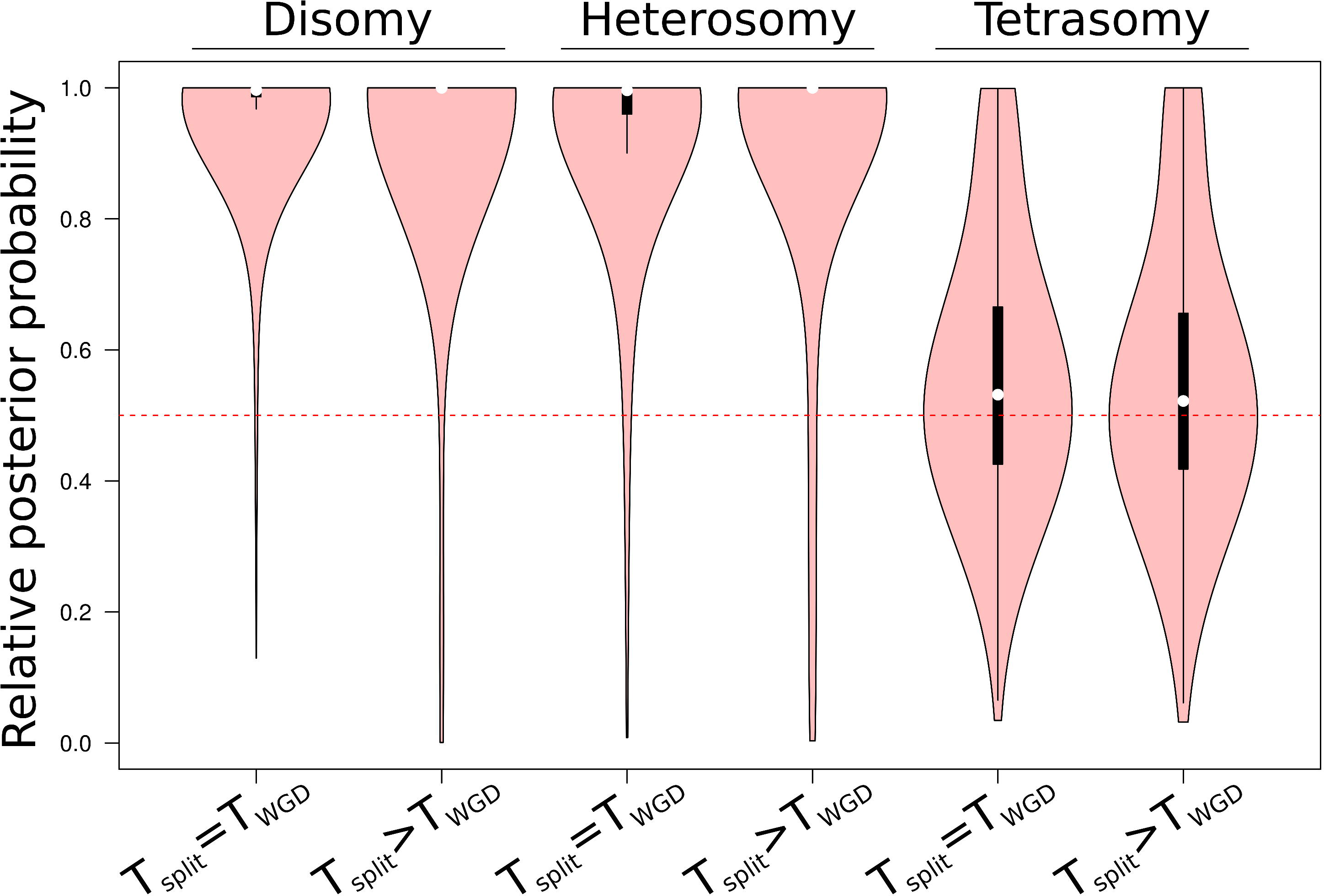
Empirical distributions of relative posterior probabilities to correctly support the co-occurrence of speciation and polyploidization regarding the mode of inheritance. Red areas represent the distributions of the estimated values of *P*(*T*_split_=*T*_WGD_ | *T*_split_=*T*_WGD_) and *P*(*T*_split_>*T*_WGD_ | *T*_split_>*T*_WGD_) obtained after model-comparisons using ABC from 1,000 pseudo-observed datasets randomly simulated when *T*_split_=*T*_WGD_ and *T*_split_>*T*_WGD_ respectively. Pairwise model-comparisons were made for disomic, heterosomic and tetrasomic mode of inheritance. The black boxes within the red areas delimit the 25% and 75% quantiles of each distribution, and the white dot show the median.

Of course, polyploidization represents one of the few realistic causes of sympatric speciation, and the coincidence of polyploidisation and speciation might be expected by default for a number of reasons; its statistical rejection is thus likely to be the more interesting outcome of ABC modeling. A newly formed polyploid population can be isolated from the parental species as a result of induced genomic incompatibilities between populations with different ploidy levels (Fowler and Levin 1984; Rodriguez 1996) and the rapid reorganization of chromosomes that result from “genomic shock” (Parisod et al. 2010). Polyploidization can also induce speciation as a result of a change in the mating system, especially the evolution of self-fertilization (Otto and Whitton 2000; Soltis et al. 2010; Pannell et al. 2004). Finally, genome doubling can extend the range of trait values, allowing the colonization of new ecological areas and leading to a geographical isolation and allopatry (Levin 1983). Cases where polyploidisation does not co-occur with speciation are likely to be rare, but testing for them using ABC modelling is straightforward.

### Inferring model parameters

We also considered the precision of ABC to retrieve parameter values for simulated models when *T*_split_=*T*_WGD_. Figure 5 shows the distributions of the ratio β = θ_i-est_/θ_i-real_,where θ_i-est_ is the median of the estimated posterior distribution of the parameter θ_i_ and θ_i-real_ is the true parameter value used to produce the pseudo-observed dataset. For the three different modes of inheritance, the distributions of β were generally centered around one, pointing to high accuracy un parameter estimation for all parameters and models. However, there were some differences in precision among the parameters and models measured by the median absolute deviation to the median of β (MAD_β_). Concerning the time of speciation (*T*_split_), the effective population sizes of the diploid (*N*_1_) and ancestral species (*N*_a_), and the proportion of genome segregating with disomic inheritance under the heterosomic model (α), ABC provides very precise and trustworthy estimates, with small MAD_β_ (Fig. 5). In contrast, the more elevated MAD_β_ measured for the effective population size of the tetraploid species (*N*_2_) suggests that we should have less confidence in this parameter estimate: caution is thus warranted when interpreting posterior distributions of *N*_2_ for all modes of inheritance when *T*_split_ = *T*_WGD_ (Fig. S1-S3).

**Figure 5:**
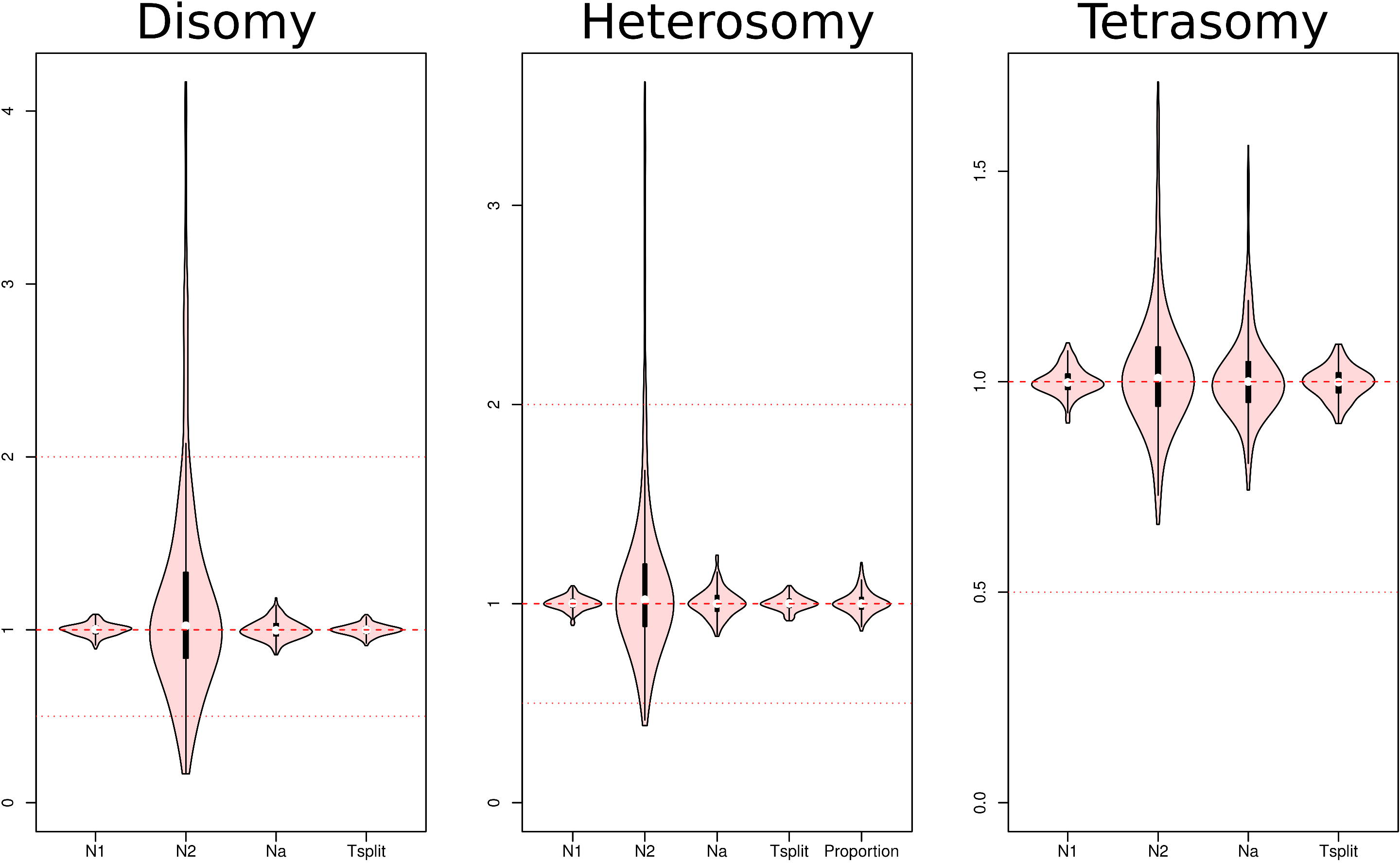
Estimates provided by ABC from pseudo-observed datasets simulated when *T*_split_=*T*_WGD_, regarding the mode of inheritance. The y-axis shows the median of the estimated posterior distribution divided by the true value for each parameter. The results are for 500 pseudo-observed datasets simulated under the disomic (Fig. S1), heterosomic (Fig. S2) and tetrasomic (Fig. S3) modes of inheritance when *T*_split_=*T*_WGD_.

Our results suggest that ABC should provide a useful tool for the joint estimation of a number of key parameters in the process of polyploidisation, such as *T*_split_, *T*_WGD_ and α, but estimates of *N*_2_ should be interpreted with caution, and estimates of the effective population size of the ancestral diploid population prior to tetraploidization (*N*_2a_) have little value. When *T*_split_ > *T*_WGD_ (Fig. 6), ABC provided precise estimates for *N*_1_, *N*_a_, α and *T*_split_. The precision achieved when estimating *T*_WGD_ and *N*_2_ were globally credible for all models, with a few caveats concerning their estimates for disomic and heterosomic autopolyploids. As might be expected, the strong bottleneck occurring at the time of polyploidization affects estimates of the effective population size of the diploid lineage before the genome duplication (*N*_2a_), as most of the informative polymorphism is lost as a result of the bottleneck. Thus, the precision achieved for estimates of *N*_2a_ was between about 8.5 (disomic autopolyploid) and 18 (heterosomic allopolyploid) times worse than that achieved for *N*_1_, as measured by MAD_β_. This lack of statistical support is evident more in terms of very wide posterior distributions than by well-defined unimodal posteriors around wrong values, i.e., it is a question of reduced precision rather than accuracy (Fig. S4–S9).

**Figure 6:**
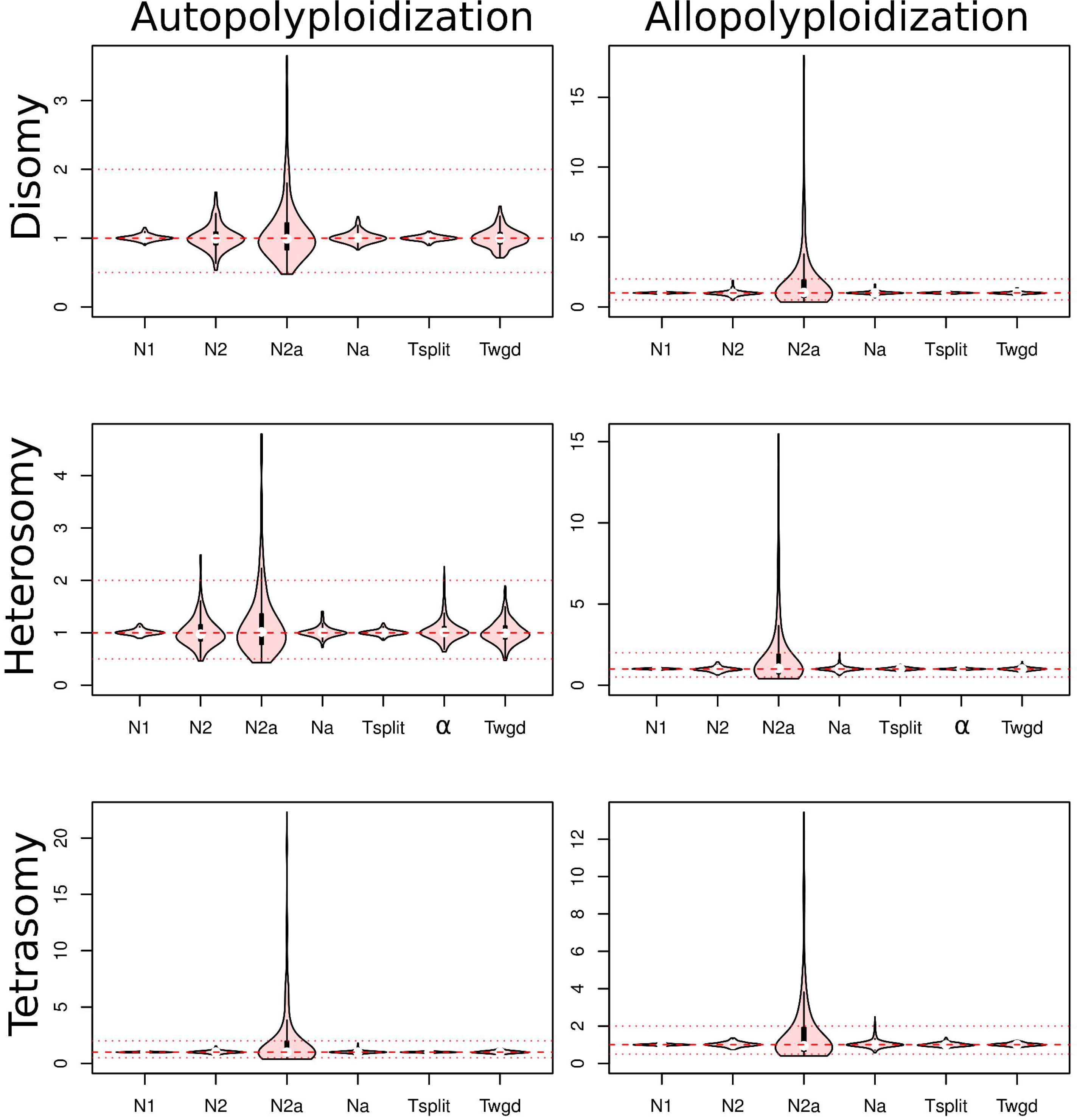
Estimates provided by ABC from pseudo-observed datasets simulated for auto- and allopolyploidization, regarding the mode of inheritance. The y-axis shows the median of the estimated posterior distribution divided by the true value for each parameter. The results are for 500 pseudo-observed datasets simulated under the disomic (Fig. S4 and Fig S7), heterosomic (Fig. S5 and Fig S8) and tetrasomic (Fig. S6 and Fig S9) modes of inheritance, and for auto- and allopolyploidization respectively.

Genome duplication has important consequences for genome structure, gene content and gene expression (Doyle et al. 2008). Inferring its date for a given biological model is therefore a key feature contributing to our understanding of molecular evolution in polyploids. In the simplest case, when polyploidization co-occurred with speciation, this date can ultimately be estimated using a molecular clock on the basis of the silent-site divergence between the polyploid species and its closely related putative diploid progenitor (Doyle and Egan 2010; Senchina et al. 2003). However, even for this simple model, the inferred date more reflects the coalescent time between diploid and polyploid lineages than the time of population subdivision (Wakeley 2009). Thus, for large ancestral population sizes, variation in coalescence depth across the genome can be large enough to complicate the use of molecular clock approaches, whereas ABC precisely provides estimate for the population separation when *T*_split_ = *T*_WGD_. If genome duplication has occurred after speciation (looking forward in time), the use of molecular divergence between duplicated genes within the tetraploid (Lynch and Conery 2000) appears to be a reasonable approach to infer the time of polyploidization, notably for allopolyploidisation. However, in addition to its weakness in dealing with tetrasomic inheritance in autopolyploids, this approach will also be highly sensitive to incomplete lineage sorting when the difference between the time of speciation and the time of polyploidization is small compared with the ancestral *N*_2a_.

### The relative efficiency of contrasting sequencing methods

We also considered how best to optimize genome sampling for model choice and parameter estimation. Overall, our results indicate that, for demographic and evolutionary inference, the number of surveyed loci is more important than having a sequenced outgroup or a larger number of sequenced individuals. With an NGS-like dataset (e.g., a large number of short contigs from an Illumina run), the absence of information from a putative outgroup had only a very small negative effect on our ability to distinguish between auto- and allopolyploidisation (Table 1), to identify correctly the mode of inheritance (Table 2), or to test for co-occurrence (or not) of *T*_split_ and *T*_WGD_ (Table 3). Precision in parameter estimates were also highly similar with or without a sequenced outgroup, either for the three *T*_split_ = *T*_WGD_ scenarios (Table 4), or for the six auto- and allopolyploid scenarios, with very little gained in terms of MAD_β_ by including an outgroup (Table 5). It would seem from our results that the effort in acquiring data from an outgroup species is unlikely to be worthwhile if the main objective is inferring the demographic history of a diploid and derived tetraploid species–though other benefits of including an outgroup will need to be evaluated, such as determining the polarity of mutations, the quantification of fixed alleles in both lineages instead of relying on the total number of fixed differences between them (Ramos-Onsins et al. 2004), the estimation of locus-specific mutation rates.

**Table 1:** Robustness of ABC for distinguishing between auto- and allopolyploidization. Numbers represent the proportions of pseudo-observed datasets supported by ABC as having an auto- or allopolyploid origin over pairwise model-comparisons. For each combination of {mode of inheritance}x{scenario}x{sequencing strategy}x{outgroup availability}, 1,000 pairwise model-comparisons have been made.

**Table 2:** Robustness of ABC for estimating the mode of inheritance. Numbers represent the proportions of pseudo-observed datasets supported as having a disomic, heterosomic or tetrasomic inheritance over model-comparisons by ABC. For each combination of {scenario}x{sequencing strategy}x{outgroup availability}x{mode of inheritance}, 1,000 pairwise model-comparisons were conducted.

**Table 3:** Robustness of ABC for testing for the co-occurrence of speciation and polyploidization. Numbers represent the proportions of pseudo-observed datasets supported by ABC as ***T*_split_=*T*_WGD_** and ***T*_split_>*T*_WGD_** over model-comparisons. For each combination of {mode of inheritance}x{scenario}x{sequencing strategy}x{outgroup}, 1,000 pairwise model-comparisons were conducted.

**Table 4:** Precision in parameter estimates under different conditions when *T*_split_=*T*_WGD_. Reported values are the median value of the distribution of β (ratio between the estimated and the true values for each parameter producing the 500 independently random pseudo-observed datasets), and its median absolute deviation to the median (MAD).

**Table 5:** Precision in parameter estimates under different conditions for cases of auto- and allopolyploidization. Reported values are the median value of the distribution of β (ratio between the estimated and the true values for each parameter producing the 500 independently random pseudo-observed datasets), and its median absolute deviation to the median (MAD).

With data that might be obtained via Sanger sequencing, we found in most cases our inferences remained robust to decreases in the number of surveyed loci. In particular, testing for auto- versus allopolyploidyzation was still reliable for a scenario of disomic inheritance, because the fixed retention of both parental genome copies represents an unambiguous signature detected by ABC (Table 1). The robustness of this model comparison decreased for increased levels of tetrasomy in genomes, regardless of the presence of an outgroup sequence (Table 1). ABC correctly distinguished between disomic and tetrasomic models of inheritance for all of the three scenarios of speciation (Table 2); when the wrong model was inferred, it tended to be in favor of heterosomy.

Sanger-like datasets were sufficient to test for the co-occurrence of speciation and polyploidization for tetraploids with disomic inheritance (Table 3), but a greater number of loci was required for species with heterosomic inheritance. As for NGS-like datasets, this test is also not conceivable when the biological model has a polysomic inheritance. Whereas reducing the number of sampled loci had only a small negative impact on model-comparisons, such datasets were not reliable for estimating key parameters in the demographic and evolutionary process. ABC is thus only poorly able to estimate *T*_split_ and *T*_WGD_ from Sanger-like datasets for scenarios where *T*_split_ = *T*_WGD_ (Table 4), or to distinguish between auto- and allopolyploidy (Table 5). In many study systems, of course, no suitable outgroup will be available, and Sanger-like datasets that have already been generated are sometimes the only sources of information. ABC can profitably be performed on these data to infer polyploid history, particularly for model comparisons, but it will be weak for parameter estimation.

### Empirical application

As a brief case study, we evalutated the use of our approach for inferring the evolutionary and demographic history of diverence between diploid *Capsella rubella* and tetraploid *C. bursa-pastoris*, for which orthologous sequences have been published (Slotte et al. 2008; St Onge et al. 2012). First we evaluated the mode of inheritance of homeologous chromosomes in tetraploid *C. bursa-pastoris* and found that, for each of the scenarios involving simultaneous polyploidy and speciation, allopolyploidization and autopolyploidization there was always strong support for a model of disomic inheritance (Table 6–a). This result confirms conclusions based on previous analysis (Slotte et al. 2006; Hurka et al. 1989; Hurka and Neuffer 1997).

**Table 6:** Relative posterior probabilities computed along the three different analyses for the *Capsella* dataset. a)Relative posterior probabilities of *C. bursa-pastoris* being disomic, heterosomic and tetrasomic, within different demographic scenarios (polyploid speciation, allopolyploidization, autopolyploidization) b) Relative posterior probabilities of *C. bursa-pastoris* being associated to different demographic scenarios within different recombinational regimes.

Second, we statistically evaluated the relative posterior probabilities of the three alternative demographic scenarios under disomic, heterosomic and tetrasomic inheritance. Here, we found that, in all three cases, a model of allopolyploidization best explains the origin of *C. bursa-pastoris* (Table 6–b). This unambiguous support of a model of allopolyploidy with disomic inheritance was also found when we compared all nine models in the same analysis, with a posterior probability ≈ 0.82 (Fig. 1). The parameters estimated for this model (Fig. 7) point to an ancestral split between the two diploid lineages that ultimately contributed to *C. bursa-pastoris* ≈ 735,371 years ago (with a higher posterior density at 95, 95HPD, estimated between 266,414 and 1,521,383 years), and that they then hybridized to form *C. bursa-pastoris* ≈ 89,199 years ago (95HPD between 34,064 and 198,738 years).

Assuming a model with constant population size since the ancestral split, the lineage leading to *C. rubella* had an effective size of ≈ 85,000 individuals (with 95HPD between 24,966 and 205,524 individuals). The estimated current population size of each sub-genomes of *C. bursa-pastoris* (≈ 355,900 individuals; 95HPD between 138,241 and 863,222) could not be compared directly to the long-term effective population size of *C. rubella*, because we infer that population size grew rapidly at a rate of 1.343 × 10^−4^. Thus, the long-term effective population size for each of the *C. bursa-pastoris* sub-genomes is better approximated by its geometric mean rather than by the current estimate, providing a value of ≈ 842 individuals over the last ≈ 90,000 years. Given that the effective population size of a diploid lineage immediately prior to whole-genome duplication cannot be faithfully estimated (Table 5, and see above), we do not provide the posterior distribution of this parameter. However, overall, a goodness-of-fit test indicates that our inferences provide a satisfactory reflection of the empirical dataset, yielding the expected distribution of summary-statistics simulated from the joint posterior distribution of the parameters under the best-supported model (Fig S10; Cornuet et al. 2010).

**Figure 7:**
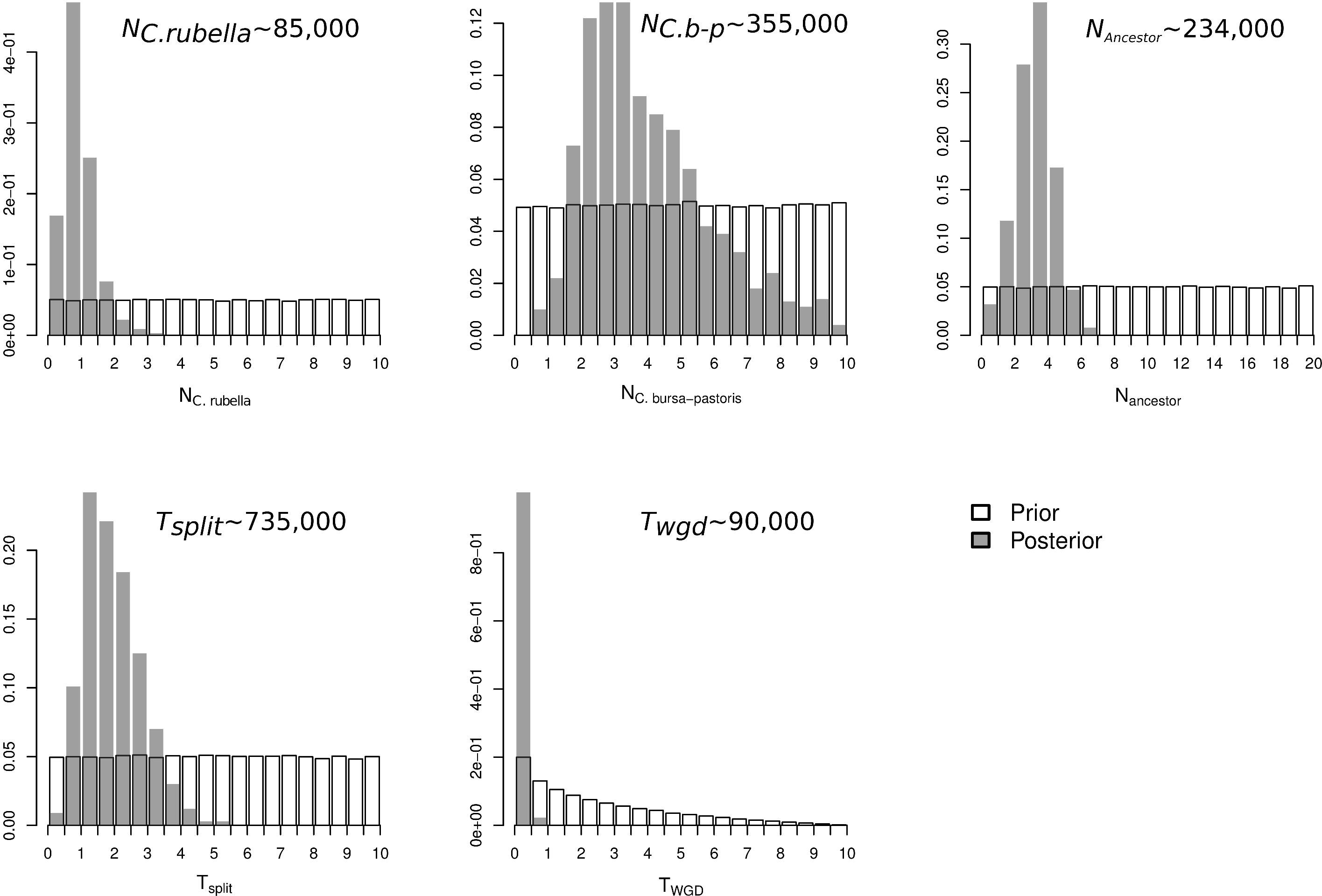
Parameter estimates of the best supported model of disomic-allopolyploidization. Prior and posterior distributions for parameters are open and shaded respectively. The median of each parameter is given along, converted into number of effective individuals or years.

### Conclusions

Our study establishes the value of ABC for inferring the evolutionary and demographic history of polyploid species on the basis of genomic data generated by state-of-the-art sequencing technologies. Not only does our approach allow us to recapture with satisfying accuracy some of the parameters used to simulate genomic data during the divergence and coalescence of lineages in the evolution of polyploid complexes; we also find that it can be applied to real data, making plausible inferences. It is particularly satisfying to note that NGS data can be used for the analyses we have tested, without any need to attribute alleles to the potentially divergent homoeologous genomes making up a polyploid lineage. Unsurprisingly, our ability to infer certain parameters was weakened for scenarios involving tetrasomic inheritance; such inheritance is analogous to effectively free gene flow between spatially isolated populations, which we should not expect to differ: tetrasomic inheritance erodes signatures of genomic subdivision and hence our ability to infer the origin of the constitutive genomes just as migration erodes our ability to attribute alleles to one deme or another. Nevertheless, it is important to note that a certain degree of disomic inheritance in a genome that displays heterosomy quickly becomes sufficient to allow accurate inference. It is also worth emphasizing that a great deal can be achieved by increasing the numbers of loci sampled, particularly for genomes displaying heterosomy. Of course, actual scenarios of polyploid are often a great deal more complex than those we have explored here, not least in complexes with higher ploidy levels and mixtures of auto- and allopolyploidy. It should, however, be straightforward to extend our approach for dealing with other more complex scenarios. Indeed, the flexibility of ABC allows the testing of quite specific models of evolution and demography. How additional model complexity in the context of polyploid evolution will affect our confidence in parameter estimation awaits further work.

## METHODS

### Models

We assessed the ability of ABC as a robust approach to investigate polyploidization by surveying a set of nine evolutionary and demographic scenarios (Fig. 1). These scenarios differ in terms of their origin of genome doubling (autopolyploid or allopolyploid), the timing of polyploidization (whole-genome duplication, WGD, is either coincident with speciation for models I, or occurred after speciation for models II), and the mode of inheritance of chromosomes (disomic, heterosomic, tetrasomic). All of the nine models describe the subdivision of an ancestral diploid population at time *T*_split_ into two populations: the first population remains diploid with a constant population size *N*_1_, while the second population experiences WGD at the time *T*_WGD_. We treated the sub-genomes of the polyploid species as two discrete populations, 2A and 2B, each sharing the same current effective population size *N*_2_. Since we assume a strong bottleneck in population size due to drastic founding effect of the new tetraploid lineage, both 2A and 2B also share the same exponential growth rate computed as −log(1/*N*_2_)/*T*_WGD_.

To simulate disomic inheritance, we considered a scenario in which there was no gene flow between 2A and 2B from time of genome duplication. As for every model, both 2A and 2B sub-genomes are also assumed to be isolated from the diploid species. We approximated the tetrasomic inheritance following Meirmans and Van Tienderen (2013) by attributing a rate of effective migration arbitrarily fixed to 10 individuals per generation, which is sufficiently high to prevent differentiation between sub-genomes (Whitlock and McCauley 1999; Wang 2004; Templeton 2006). It is worth noting that this approximation only holds up because we assumed a single panmictic tetraploid population that receives no migrants from unsampled populations. Heterosomy is implemented in our simulations as a binomial distribution of size equal to the number of investigated loci, and a probability for a given locus to exchange alleles between sub-genomes equal to α. Thus, for heterosomic models, α represents the expected proportion of the genome experiencing tetrasomic inheritance, and (1-α) represents the expected proportion of the genome experiencing disomic inheritance.

In allopolyploid models, the diploid species and the 2-A lineage first come together at time *T*_WGD_, following which both of the ancestral diploid lineage and the 2-B lineage join at time *T*_split_. In autopolyploid models, the first fusion backward in time occurred between 2-A and 2-B at time *T*_WGD_, then the ancestral tetraploid lineage with effective population size *N*_2-anc_ is joined to the first lineage at time *T*_split_. Finally, *T*_split_=*T*_WGD_ for the three models I, and *T*_split_>*T*_WGD_ for the six models II.

### Coalescent simulations

For each model, we obtained a set of 5 × 10^6^ multilocus simulated data sets using msnsam (Ross-Ibarra et al. 2008), a modified version of the ms software (Hudson 2002), allowing for variation in sample size among loci under an infinite-site mutation model. The command lines used to simulate the different models are shown in supplemental text S1. We used the software Priorgen to generate the prior distributions given the nine models, and mscalc to compute the summary statistics described below from the output of msnsam (Ross-Ibarra et al. 2008, 2009; Roux et al. 2011, 2013).

We used a wide uniform prior distribution [0-10] common to all models and shared by all parameters in order to consistently explore a large panel of realistic biological scenarios. Values for α have been randomly sampled in the uniform prior [0.025-0.975]. To convert coalescent units in natural demographic units, effective population sizes *N*_A_, *N*_1_, *N*_2-anc_ and *N*_2-A_ (or N_2-B_) have to be multiplied by the effective population size of the virtual reference population arbitrarily fixed to 10^5^. For example, following the used prior for *N*_1_, this parameter can take a value between zero and one million. *T*_split_ and *T*_WGD_ have to be multiplied by 4.10^5^ to obtain the number of generations in demographic units, so that *T*_split_ was sampled between zero and four million generations.

For each simulated dataset, we computed an array of summary statistics related to polymorphism and divergence, as is widely used in the literature (Wakeley and Hey 1997; Becquet and Przeworski 2007; Ross-Ibarra et al. 2008, 2009; Roux et al. 2011, 2013;Fagundes et al. 2007): Tajima’s π (Tajima 1983), Watterson’s *θ*_W_ (Watterson 1975), *F*_ST_ estimated as 1- π_S_/ π_T_ (where π_S_ is the mean pairwise diversity for both species and π_T_ is the pairwise diversity calculated from the total alignment), Tajima’s *D* (Tajima 1989), total interspecific divergence *Div*, and the net interspecific divergence *netDiv* (=*Div*-π_S_). We add two alternative sets of summary statistics depending on the availability of a sequenced outgroup for the user. In cases where a sequenced outgroup was assumed to be available to orientate mutations (Ramos-Onsins et al. 2004), we also used as summary statistics the number of derived biallelic positions exclusively polymorphic in the diploid species (*SxA*) and the number of derived positions exclusively polymorphic in the tetraploid species (*SxB*). Concerning the latter category, a sequenced tetraploid individual is seen as the merging of two copies from sub-genome 2-A and two copies from sub-genome 2-B without the possibility of distinguishing the two. We counted the number of polymorphic sites shared by both species (*Ss*), *i.e.*, a biallelic polymorphism found in the diploid species as well as in the merged 2-A and 2-B tetraploid species. We computed the number of positions for which a derived allele is fixed in the first species and the ancestral allele is fixed in the second one (*SfA*). In the same way, we defined a *SfB* site as a genomic position for which an ancestral allele is fixed in the diploid species and the derived allele is fixed in both 2-A and 2-B subgenomes. The position *SfAxB* (or *SfBxA*) defines a derived allele fixed in the diploid species (or in the tetraploid), but still segregating in the tetraploid species (or in the diploid). In the situation where a sequenced outgroup was assumed not to be available to the investigator, we computed *SxA* as the total number of polymorphic positions specific to the diploid species regardless of the fixed allele in the tetraploid species, and used the same rational to compute *SxB*. In this situation with no outgroup sequence, *Sf* is equal to the sum of *SfA* and *SfB*. Finally, we used the mean and the standard deviation of each quantities computed over all of the loci as summary statistics.

### Alternative sequencing strategies

We used coalescent simulations to assess the robustness of ABC through four alternative experimental schemes: datasets with or without an outgroup; and data generated with NGS-like or Sanger-like sequencing. While the presence or absence of an outgroup only impacts the summary statistics described above, the sequencing strategy mainly impacts the sampling size as well as the number of re-sequenced loci. For the Sanger-like dataset, we assumed the availability of a set of 30 loci of mean synonymous length equal to 110 (SD = 10) resequenced for around 60 sampled chromosomes (SD = 10); this is similar to Sanger-like datasets recently analysed in the literature (Roux et al. 2011). For the NGS-like dataset, we assumed a set of 852 loci of mean synonymous length equal to 70 (SD = 30) resequenced for around 16 sampled chromosomes (SD = 4); this is similar to the dataset investigated in Roux et al. (2013). For each of the four experimental strategies, we simulated 1,000 pseudo observed datasets under the nine scenarios investigated.

### ABC analysis

#### Model selection

For each model comparison, we evaluated the relative posterior probabilities of the alternative models using a feed-forward neural network, implementing a nonlinear multivariate regression by considering the model itself as an additional parameter to be inferred with the R package “abc” (Csillery et al. 2012). The 0.025% replicate simulations nearest to the observed values for the summary statistics were selected, and these were weighted by an Epanechnikov kernel that reaches a maximum when *S*_obs_=*S*_sim_, where *S*_obs_ are the summary statistics computed from the observed or the pseudo observed datasets. Computations were performed using 50 trained neural networks and 10 hidden networks in the regression.

### Parameter estimation

Posterior distributions of the parameters describing the best model were estimated using a nonlinear regression procedure. Parameters were first transformed according to a log-tangent transformation (Hamilton et al. 2005). Only the 2,000 replicate simulations that provided the smallest associated Euclidean distance *δ* = ||*S*_obs_ **-** *S*_sim_|| were considered. The joint posterior parameter distribution was then obtained by means of weighted nonlinear multivariate regressions of the parameters on the summary statistics. For each regression, 50 feed-forward neural networks and 20 hidden networks were trained using the R package “abc”.

### Capsella dataset

We downloaded published sequences obtained in *C. rubella* and *C. bursa-pastoris* (St Onge et al. 2012; Slotte et al. 2008) from GenBank (accession numbers EF683687–EF684898 and JQ418636–JQ419488), making a total of 13 aligned loci. Only bi-allelic polymorphism at silent sites (introns + synonymous positions from exons) were considered in the analysis, based on the GenBank annotations. For each of the nine alternative models, we ran three million multilocus simulations, with the following uniform prior distributions. *N_C. rubella_* [0-1,000,000]; *N_C. bursa-pastoris_* [0-1,000,000]; *N_ancestor_* [0-2,000,000]; *T_split_* [0-4,000,000]; *T_WGD_* [0-*T_split_*]. A mutation rate of 1.5 × 10^−8^ per site and per generation was then used to rescale the coalescent units in demographic units, assuming a generation time of one year. We finally applied the same ABC procedure as described above. Simulations and ABC analysis for the *Capsella* dataset were run for a total of three hours when parallelized on a cluster using 150 CPUs.

## DATA ACCESS

The used code is available at: http://www.unil.ch/webdav/site/dee/shared/softs/ABC_WGD.tar.gz

## ACKNOWLEDGMENTS

We are grateful to Jonathan Wendel for suggestions and comments the manuscript. The computations were performed at the Vital-IT Center for high-performance computing of the SIB Swiss Institute of Bioinformatics (http://www.vital-it.ch). CM was supported by a postdoctoral fellowship of the Swiss Nationan Science Foundation awarded to JRP.

---

**Figure S1: Comparison between posterior distribution and real value for the disomic inheritance when *T*_split_=*T*_WGD_ for 500 pseudo-observed datasets.**

For each parameter, the red vertical line indicates the real parameter value used to simulate the pseudo-observed dataset, and the histogram shows the estimated posterior distribution obtained using ABC.

**Figure S2: Comparison between posterior distribution and real value for the heterosomic inheritance when *T*_split_=*T*_WGD_ for 500 pseudo-observed datasets.**

For each parameter, the red vertical line indicates the real parameter value used to simulate the pseudo-observed dataset, and the histogram shows the estimated posterior distribution obtained using ABC.

**Figure S3: Comparison between posterior distribution and real value for the tetrasomic inheritance when *T*_split_=*T*_WGD_ for 500 pseudo-observed datasets.**

For each parameter, the red vertical line indicates the real parameter value used to simulate the pseudo-observed dataset, and the histogram shows the estimated posterior distribution obtained using ABC.

**Figure S4: Comparison between posterior distribution and real value in cases of autopolyploidization with a disomic inheritance, over 500 pseudo-observed datasets.**

For each parameter, the red vertical line indicates the real parameter value used to simulate the pseudo-observed dataset, and the histogram shows the estimated posterior distribution obtained using ABC.

**Figure S5: Comparison between posterior distribution and real value in cases of autopolyploidization with a heterosomic inheritance, over 500 pseudo-observed datasets.**

For each parameter, the red vertical line indicates the real parameter value used to simulate the pseudo-observed dataset, and the histogram shows the estimated posterior distribution obtained using ABC.

**Figure S6: Comparison between posterior distribution and real value in cases of autopolyploidization with a tetrasomic inheritance, over 500 pseudo-observed datasets.**

For each parameter, the red vertical line indicates the real parameter value used to simulate the pseudo-observed dataset, and the histogram shows the estimated posterior distribution obtained using ABC.

**Figure S7: Comparison between posterior distribution and real value in cases of allopolyploidization with a disomic inheritance, over 500 pseudo-observed datasets.**

For each parameter, the red vertical line indicates the real parameter value used to simulate the pseudo-observed dataset, and the histogram shows the estimated posterior distribution obtained using ABC.

**Figure S8: Comparison between posterior distribution and real value in cases of allopolyploidization with a heterosomic inheritance, over 500 pseudo-observed datasets.**

For each parameter, the red vertical line indicates the real parameter value used to simulate the pseudo-observed dataset, and the histogram shows the estimated posterior distribution obtained using ABC.

**Figure S9: Comparison between posterior distribution and real value in cases of allopolyploidization with a tetrasomic inheritance, over 500 pseudo-observed datasets.**

For each parameter, the red vertical line indicates the real parameter value used to simulate the pseudo-observed dataset, and the histogram shows the estimated posterior distribution obtained using ABC.

**Figure S10: Principal component analysis of summary statistics simulated from the posterior distribution under the best-supported model.**

A principal component analysis is performed in the space of summary statistics after 10,000 simulated data sets using the posterior distribution under the allopolyploid scenario with disomic inheritance (blue dots).

The observed *Capsella* target is added on each plane of the PCA (red dot).

